# Connectal Coding: Discovering the Structures Linking Cognitive Phenotypes to Individual Histories

**DOI:** 10.1101/610501

**Authors:** Joshua T. Vogelstein, Eric W. Bridgeford, Benjamin D. Pedigo, Jaewon Chung, Keith Levin, Brett Mensh, Carey E. Priebe

## Abstract

Cognitive phenotypes characterize our memories, beliefs, skills, and preferences, and arise from our ancestral, developmental, and experiential histories. These histories are written into our brain structure through the building and modification of various brain circuits. Connectal coding, by way of analogy with neural coding, is the art, study, and practice of identifying the network structures that link cognitive phenomena to individual histories. We propose a formal statistical framework for connectal coding and demonstrate its utility in several applications spanning experimental modalities and phylogeny.

---

Scientific explanations are characterized by modeling mechanisms of phenomena that describe (i) the constituent parts, (ii) the properties of those parts, and (iii) the interactions among them [1]. What we measure and model is often limited by technology. In brain sciences, 20^th^ century innovations enabled studying the parts and their properties, while studying interactions was limited [2] and laborious [3]. For example, nanoscale electron microscopy (EM) enabled studies of ultrastructure [4], microscale light microscopy and physiology enabled studies of single cells [5], and macroscale magnetic resonance imaging (MRI) enabling studies of brain regions [6,7]. 21^st^ century innovations include serial EM for measuring subcellular interactions [8], improved LM [9–14], and MRI [15–18] to estimate interactions within and across brain regions.

These 21^st^ technologies now enable modeling brain connectivity, in a complementary fashion to models of brain activity that were developed in the 20th century [19,20]. Models of brain *activity*, typically referred to as *neural coding*, link patterns of brain activity to past, ongoing, and future events. By contrast, models of brain *connectivity*, which we refer to as *connectal coding*, link patterns of brain connectivity to past, ongoing, and future events. The nature of the patterns, events, and links change by virtue of switching focus from activity to connectivity. Moreover, the statistical models one can leverage to learn those links from the data must also change.

The goal of this manuscript is to introduce in clear terms, motivate from first principles, and formalize this emerging approach to studying the brain. While neural activity coding is well established and widely accepted as a (possibly *the*) legitimate framework for studying the brain, connectal coding remains in its infancy. Below we outline our rationale for why connectal codes are not just valuable, but required to unify 20th and 21st century mentalities to model the parts, their properties, and interactions among them together to infer improved scientific explanations of cognitive phenomena.

## Modeling Brains as Networks

The idea of the brain as a network dates back to the late 19th century, with Ramon y Cajal’s discovery of dendritic spines using Golgi’s stain. This marked the founding of the “neuron doctrine”, for the first time asserting that the brain is not, in fact, a syncytium [5]. In 1943, McCulloch and Pitts wrote down a mathematical model of the brain as a network, and proved that such networks could be “Turing-complete” (that is, they can solve any computational problem) [21], thereby founding the field of artificial intelligence [22]. Shortly thereafter, Hebb introduced “cell assemblies” as essentially networks of neurons that are jointly active [23]. A few decades later Little [24], and then Hopfield [25], introduced Little-Hopfield networks (which were the first recurrent neural networks and led to the founding of connectionism [26]). Little and Hopfield leveraged an idea from statistical physics known as the Ising model [27], positing that these assemblies could be used to store information. But in the entire 20th century, it was rare to use the tools of “graph theory” to study brain networks, even though graph theory was founded by Leonhard Euler in 1736 [28].

This all changed with the introduction of the term “connectome” by Sporns *et al*. and Hagmann in 2005 [15,16]. Since then, >3,600 papers have been published using the term. Because many of those papers define connectome differently, we provide our definition below:

### Definition

A *connectome* is an abstract mathematical model of brain structure, denoted *G*, and is a set of two kinds of objects:

1. Vertices (or nodes), *V*: A vertex represents a biophysical entity of the brain. In a connectome, one defines a set of constraints, including the spatial extent (e.g., the left mushroom body, a cortical column, or the whole brain), spatial resolution (e.g., a cell, cellular compartment, or a cellular ensemble), type (e.g., a neural, glial, or perivascular cell), and developmental stage (e.g., postnatal day six). The nodes of a connectome are all the nodes satisfying those constraints.
2. Edges (or links), E: An edge between any pair of nodes represents the presence (and lack of edge represents the absence) of a connection or communication between those nodes. In a connectome, that connection/communication must satisfy another set of constraints, including the kind of communication (e.g. transmission of electrical charges, neurotransmitters, physical opposition, or fiber bundles) and the temporal duration under which these communications may be present (e.g., during a particular developmental phase, or during a traumatic injury including brain compression, etc.). The edges of a connectome are all the edges satisfying these constraints between the above described set of nodes. Note that implicit in this definition is that the connections for which edges count in connectomes are direct.

Under this simplest definition of what constitutes a connectome, it is common to represent the connectome via a two-dimensional (2D) array, *A* (Figure 1). In this representation, each row/column pair corresponds to a node, and edges between a pair of nodes *u* and *v* are depicted by a non-zero entry in the corresponding element of the array, i.e., *A(u,v)=1*. It is tempting to think of this as a matrix, and it certainly is from the computer science perspective. But it is decidedly *not* a matrix from the mathematics perspective, where matrices are linear operators, e.g. *y* = *Ax*. Whereas matrix algebra can be applied to study these representations of graphs, we often find it helpful to keep in mind that these “adjacency matrices” are special kinds of 2D arrays. The row identities are inextricably linked to column identities, a property that is not generally true for arbitrary matrices, and changes the kinds of statistical procedures appropriate for these data.

**Figure 1:**
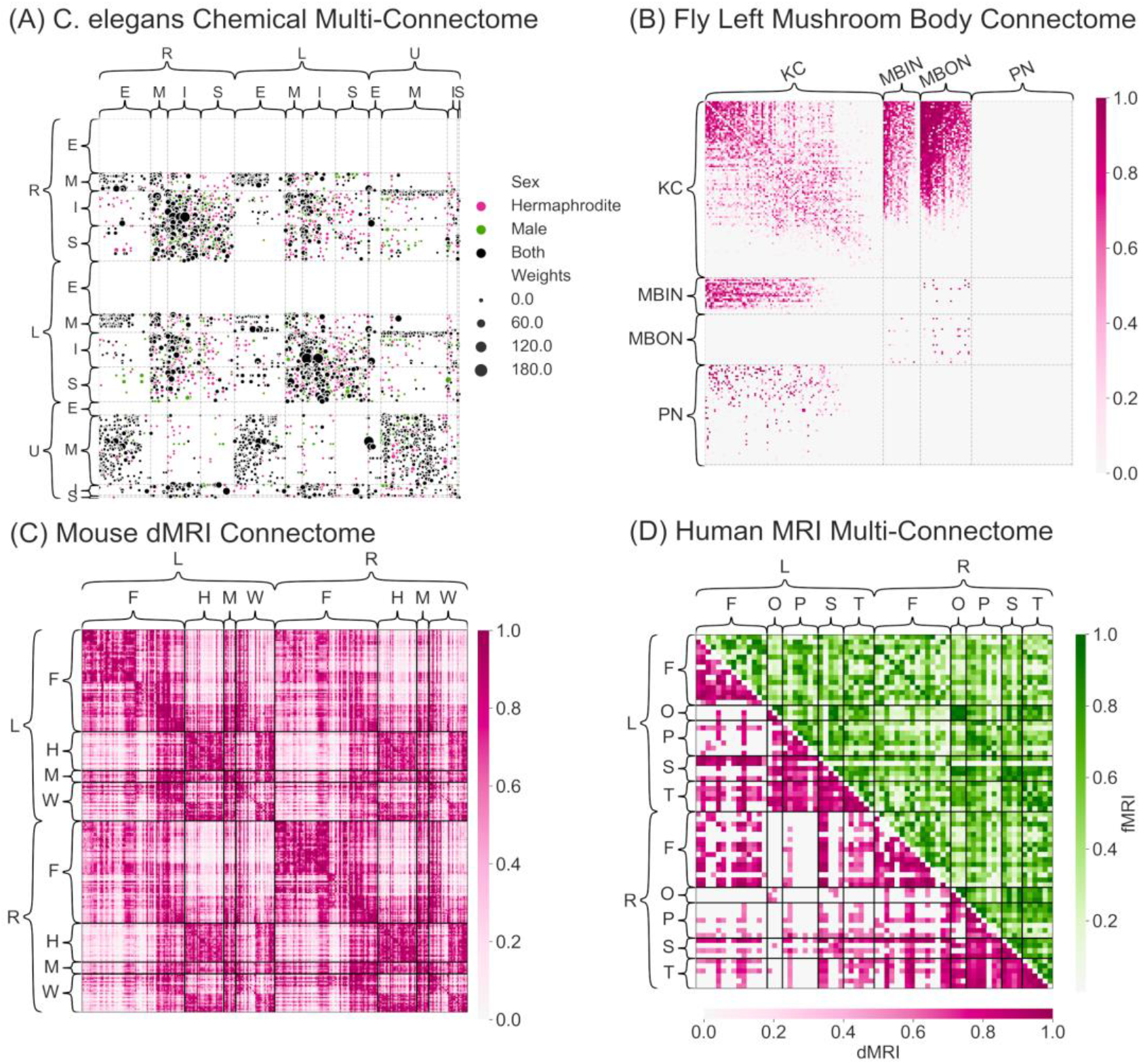
Estimated connectomes spanning four levels of the phylogenetic tree, each estimated using different experimental modalities and spatial resolutions, ranging from nanoscale (electron microscopy) to macroscale (MRI regions). **(A)** C. elegans chemical multi-connectome estimated from male and hermaphrodite connectomes. Size of circles corresponds to the number of synapses between two neurons. Multiscale node labels: left (L) and right (R), which are bilateral pairs, as well as unpaired (U); and four types per side: endorgans (E), interneurons (I), motor neurons (M), and sensory neurons (S). **(B)** Left mushroom body connectome estimated from Drosophila melanogaster. Nodes represent neurons, and are assigned into kenyon cells (K), input neurons (I), output neurons (O), and projection neurons (P). Color intensity corresponds to the number of synapses between two neurons. **(C)** Mouse estimated connectomes obtained from dMRI scans. Nodes represent regions of the brain, and are assigned into right (R) and left (L) hemispheres and then further assigned into superstructures such as frontal (F), hindbrain (H), midbrain (M), and white matter (W). Color intensity corresponds to degree of connectivity between two regions. (D) Human multi-connectome estimated from averaging 3,067 dMRI and 1,760 fMRI human connectomes. Since both networks are undirected, only the upper triangle of fMRI connectome and lower triangle of dMRI connectome is shown. Nodes represent regions of the brain, and are assigned into right (R) and left (L) hemispheres and then further assigned into frontal (F), occipital (O), parietal (P), and temporal (T), as well as subcortical structures (S). Color intensity corresponds to degree of connectivity and correlation for dMRI and fMRI, respectively, between two regions.

Moreover, in classical graph theory, the graph is simply the tuple *G=(V,E)*, and lacks additional structure. For connectomes to serve as a model for brain structure that can link to past, ongoing, and future events, they often require some additional structure. The most common additional structure is *edge weights*, such that each binary edge is associated with a magnitude that can take any continuous value. More complicated and nuanced edge attributes abound in connectomics. For example, when edges correspond to synapses, they might be attributed with weights, locations, directions of transmission, neurotransmitters, etc. Similarly, for connectomes, it is natural for nodes to have attributes. The most common attribute of a node is a semantic label, such as the Mauthner neuron [29], or Purkinje cell, or primary visual cortex. Nodes in connectomics, like edges, can have other attributes as well, such as location, volume, and shape. Finally, when studying populations of connectomes, the entire graph may be endowed with attributes, such as a weight.

The above definition clarifies what a connectome is for this manuscript, but not what it is not. It is not many things; we illustrate a few. The shape, size, or morphology of a set of brain regions or neurons does not comprise a connectome, nor does the set of all spike trains of a brain. These features can be attributes of nodes, but without also characterizing the edges, one has not modeled a connectome. Similarly, correlation between a pair of nodes cannot be defined as an edge (though it may be used to estimate the presence of an edge). This is because connectomes, as defined above, are models of brain structure and anatomy, and correlation is an emergent property of dynamics on that model, rather than a description of the model itself. While many connectomics papers disagree with this definition, we find it useful, and use it for the remainder of the manuscript.

An implication of this definition is that one could simultaneously model a given brain with many different connectomes at different times, or at different resolutions, or of different types, etc. Moreover, the neuron-level model of brain structure does not have an elevated status over, say, compartments of neurons, glia, or brain regions; rather, each scale and type of node can serve as a perfectly adequate model of part of the brain, and those parts may turn out to be the most important parts to explain any particular past or future event.

Given all this, one can measure parts of a brain to estimate a connectome from many different experimental modalities, each of which has its shortcomings. At cellular resolution, as early as 2001, several papers began characterizing how to estimate connectomes (without using the word connectome) from physiology data [30–33]. More recent developments in cellular resolution connectome estimation incorporated unobserved variables [34,35], called “confounders” in the causal inference field [36]. At the millimeter scale, diffusion magnetic resonance imaging data (dMRI) is known to exhibit both false positives and false negatives, as compared to gold standard methods [37]. By the same token, one can use functional MRI (fMRI) data to estimate edges. The most common approach by far is to simply use correlations [17]; these approaches are also known to yield problematic estimates for a number of reasons, including susceptibility to various exogenous variables, such as time of day, week, and month [38]. Methods for addressing confounders have also been proposed in the fMRI literature [39], and while widely cited, are still largely underappreciated.

## Example Estimated Connectomes

We show several different previously published estimated connectomes from different species spanning the phylogenetic tree, with widely disparate definitions of what constitutes nodes and edges (Fig. 1). There are many possible ways to visualize a connectome [40–42]; we choose a relatively simple way, adjacency matrices, as described above. We indicate the strength of connection as either the size or contrast of the corresponding matrix element. Showing an adjacency matrix requires first sorting all the nodes in some order; we choose to sort by region or type, and within that by degree (total weight of connections per node), but other sortings could be equally informative. For certain connectomes, experimentalists have measured multiple types of edges. We show these “multi-connectomes” with different colors for the different edge types.

A. **Caenorhabditis elegans (worm).** C. elegans is the only animal for which we have estimated a complete connectome where the nodes represent neurons [3,43,44]. That is, every neuron in the animal is a node, and every edge has been estimated. This connectome has two types of edges corresponding to two types of neural activities: (1) chemical synapses for neurochemical release (Fig. 1A), and (2) gap junctions for electrical activity. Each edge’s strength (or weight) corresponds to the approximate total volume of synapses between its parent neurons. C. elegans has two sexes, male and hermaphrodite, with different numbers of neurons (hermaphrodite 302, male 385, with 290 overlapping shown in Fig. 1A) [45,46]. These connectome estimates are derived by cumbersome manual tracing of axons and dendrites and identification of synapses, in nanoscale electron micrographs, updated by Varshney et al. [47], Bentley [48], and most recently by Cook et al. [49].
B. **Drosophila melanogaster (fly).** Eichler et al. [50] published an estimate larval Drosophila connectome of the left (Fig. 1B) and right mushroom body, also derived from serial electron microscopy, using only chemical synapses. These edges are weighted (based on counting the number of synapses between a pair of neurons), and directed (meaning connections can be from *u* to *v* and not vice versa) [51].
C. **Mus musculus (mouse).** Calabrese et al. [52] generated a high-resolution connectivity estimate using ex vivo diffusion magnetic resonance imaging (dMRI). This network is undirected, as dMRI lacks directional information, and weights correspond to the number of tracts estimated to go between regions. Because we do not know the mapping from the absolute magnitude of connection weights to any physical connection, we rescale these weights to be between zero and one, and depict them on a log scale (Fig. 1C).
D. **Homo sapiens (human).** The Consortium for Reliability and Reproducibility collects multiple measurements of functional resting-state, anatomical, and/or diffusion magnetic resonance imaging (MRI) per individual [53]. The functional estimated connectomes are Pearson correlation matrices, converted to ranks and then normalized between zero and one, because such a representation is more reliable than raw or thresholded correlations [54]. The diffusion estimated connectomes are normalized as described above for the mouse. This multi-connectome estimate is derived from averaging the entire dataset of 3,067 diffusion and 1,760 functional MRI connectomes [53] (Fig. 1D).

## The Purpose of Brain Codes

A code is a (potentially stochastic) system of rules to convert information from one representation into another [55,56].^1^ For example, neural activity coding can be thought of as converting information from past and ongoing events (stimuli and behavior) to neural activity, and from neural activity to future events (predictions and behaviors). In other words, neural activity codes correspond to the brain’s *representation* of information. In contrast, connectal coding can be thought of as converting information from past and ongoing events (ancestral, developmental, and experiential) to brain connectivity, and from brain connectivity to future events (behavioral tendencies). In other words, connectal codes correspond to the brain’s *storage* of information. Therefore, brain activity and connectal codes serve complementary roles in understanding the relationship between genetics, body, world, and brain.

An important aspect of both activity and connectal codes is that they are stochastic. In other words, a given ongoing stimuli/behavior can stochastically manifest in multiple different patterns of activity, and a given past event can stochastically manifest in multiple different patterns of connectivity. The inverse is also true: activity/connectivity can represent multiple different current/past/future events, respectively. This stochastic property of brain codes is actually required for operating in the world. Even human brains, given finite physical and energetic resources, are incapable of representing and storing the large amount of information impinging on our sensory faculties. In a similar fashion, the connectivity of a human brain is too complex to be explicitly prescribed by their genome. To be concrete, consider that a human brain contains approximately 10^11^ neurons [58] and 10^15^ connections between pairs of neurons, and yet we only have about 10^4^ genes [59]. Thus, for the genome to encode every single synapse would require five to six variants per gene, and literally every variant of every gene would encode the brain’s synapses. More likely, the genome encodes the “blueprint”, that is, a number of statistical principles governing the probability of connections between nodes across development, as well as all the rules for learning new connections due to activity-dependent plasticity. Those rules are the principles of connectal coding.

## The Role of Connectomes in Connectal Coding

The above definition of connectome sets the stage for understanding the relationship between connectomes and other aspects of an individual or population. By way of analogy, recall that a genome is the complete genetic sequence of an individual, whereas a *genotype* is the part of the genetic sequence of an individual that associated with a particular phenotype. In that sense, a *connectotype* is the collection of nodes and edges (and potentially their attributes) associated with a given phenotype. We consider two kinds of phenotypes here: *individual histories* and *cognitive phenotypes*. By individual histories, we mean ancestral, developmental, and experiential histories; we may desire to understand the relationship between connectome and genome, connectome and developmental stage, or connectome and experience. By cognitive phenotype we mean a set of observable characteristics of an individual related to their cognition, including personality traits, memories, beliefs, skills, preferences, and psychiatric or learning disorders. We may desire to understand the relationship between connectome and these phenotypes as well. Regardless of which particular kind of phenotype, there may exist one (or many) connectotypes associated with it. Connectal coding is the study of brain structures that encode that information.

Like genotypes, there is not a one-to-one mapping between connectotypes and phenotypes, rather, a given connectotype could stochastically encode in different cognitive phenotypes at different times, and a given cognitive phenotype could be associated with many different connectotypes. For example, the stomatogastric ganglion circuit of a crab can exhibit similar network activity from disparate circuit parameters [60].

In light of this, our view on connectomics is that its primary value is in *generating hypotheses* about connectotypes. This is in contrast to the relatively low-throughput, more classical approach to studying neural circuits. For example, detailed physiological and anatomical characterization of the sound localization circuit of a barn owl [61] effectively confirmed models that explained a certain phenotype. However, the process of hypothesis generation was arduous, taking decades, including numerous models that, in retrospect, were not possible given the underlying neuroanatomy. If one could rapidly collect data about many interwoven neural circuits, and then use those data to screen and/or filter various circuits as implicated (or not) in a given behavior, then subsequent experiments could further refine those results. Thus, connectomes may not on their own provide data sufficient to *test* hypotheses about how certain genotypes are linked to certain connectotypes and/or how certain connectotypes are linked to certain phenotypes. Nonetheless, if these data can accelerate the hypothesis-generation process to hone in on a small set of plausible models, they will be extremely valuable.

Of note, in connectal coding, the role of estimating connectomes is not about describing the basic anatomical properties of connectomes, or modeling connectomes as an end unto themselves. Rather, in connectal coding, connectomes are interesting insofar as they participate in the understanding of the relationship between brain structure and individual histories or cognitive phenotypes. Part of the rationale of this focus is that most current experimental approaches for estimating connectomes are so error prone, that estimates of the statistical properties of brain networks are difficult to interpret in terms of the underlying biology. In fact, even if the part of the network that is observed is largely correct, if it represents just a subsample of the network of interest, then the resulting network features can be quite different from those features of the entire network [62,63]. Regardless, building statistical model relating connectypes to phenotypes requires models of connectomes.

## Models of Connectomes

Every connectomics study utilizes some mathematical and statistical approach to support the scientific claims. We organize those approaches into three categories, and demonstrate that only one of these frameworks is sufficient for connectal coding, although all three provide complementary insights and perspectives on the connectomes themselves.

The most common approach, dubbed the “bag of edges” (or edgewise statistics) framework. [64,65], treats each edge independently, without taking into account interactions or relationships between them. Such univariate approaches allow researchers to identify easily interpretable relationships between phenotypes and edge weights. However, this approach requires performing many statistical tests, which must be corrected for multiple comparisons to adequately control for the number of false positives. Standard correction techniques such as false discovery rate [66] do not model the dependencies between edges, and therefore may result in overly liberal or conservative corrections [67]. Alternate correction techniques such as network-based statistics [68] or group Benjamini-Hochberg corrections [69] leverage information about the group structure of connectomes to increase statistical power, while attempting to control false positives. Network-based statistics, however, lacks theoretical guarantees identifying the settings in which it successfully controls false positives. Without such an understanding, interpreting its results is problematic. Benjamini-Hochberg has strong theoretical guarantees, but in models that are inappropriate for connectomics data, in that their assumptions are often grossly violated (and rarely checked). Bonferroni corrections are widely believed to be overly conservative [70] and therefore lack the sensitivity desired for connectal coding.

A second popular approach we dub the “bag of features” framework. In this approach, multiple graph-wise or node-wise statistics are calculated [71] and compared. Possible features include degree distribution, degree sequence, clustering coefficient, number of triangles or other motifs, small worldness, efficiency, and modularity [72]. While computing these statistics can be informative about the properties of a given connectome, using them as features to explain differences between genotypes or phenotypes faces serious drawbacks. First, for any particular feature, vastly different networks can produce the same value [73]. Second, for a connectome with *n* nodes, there are *2^n^*^n^* possible subgraphs, each of which could reasonably be considered a feature. Therefore, one cannot reasonably search all features (as *2^n^*^n^* is larger than the number of atoms in the universe for *n* > sixteen!). It is therefore unclear (and somewhat arbitrary) how one should choose relevant features for a particular dataset. Third, and perhaps most problematic, is that the different features are not typically independent of one another. Therefore, if the question is whether a given phenotype depends on a particular feature being a specific value, it is impossible to determine whether *that* particular feature is responsible. Rather, the phenotype may be dependent upon a subset of the exponentially many features that correlate with both the feature of interest and the phenotype. No experiment could test whether a feature is uniquely informative with regard to the covariate of interest, even in theory. Thus, studying most network features will fail to yield the principles of connectal coding.

The third framework, which is much less popular in connectomics, but much more popular in network statistics, is statistical modeling of networks [74,75]. The key conceptual herdle associated with using statistical models of networks to model connectomes is that it is a model of the entire network, rather than just the edges or features, as a random variable. This network is a complex high-dimensional random variable, with built-in structure and relationships. The vast majority of work in network modeling focuses on modeling a single network, typically undirected, unweighted, and lacking in network-wise, node-wise, or edge-wise attributes [76]. However, connectomes are typically weighted, sometimes directed, and always include at a minimum node-wise labels (e.g., which cell, cellular compartment, or cell ensemble corresponds to a given node). Moreover, understanding the relationship between connectypes and phenotypes typically requires comparing multiple connectomes. Much of the existing work comparing multiple networks ignores the unique node labels that are often available in connectomics [77]. Therefore, a full accounting of connectotypes will require statistical models of populations of networks with complex attributes. Although a comprehensive theory remains absent, we build connectal coding on the foundational work of network modeling [78].

### Statistical Models of Connectomes

The simplest random network (graph) model is the Erdos-Renyi model, in which each edge is sampled identically and independently [79]. This binary model is the connectal coding homolog to the Poisson process in neural coding, which asserts that a neuron’s spikes are sampled identically and independently [20]. Although these models are too simple to explain much, they are excellent starting grounds to build more complex models, such as the inhomogeneous Erdos-Renyi model, which assumes that each edge is an independent coin flip, but each edge can be sampled with a different probability [76]. The next simplest binary models are stochastic block models (SBM) [80–83]. In an SBM, there are *K* groups, and each node is in one group. Each edge is again sampled independently, but the probability of a connection between a pair of nodes is no longer equal everywhere; rather, it is determined by each node’s group (homologous to a heterogeneous Poisson process, where spiking has *K* different rates, dependent on which of *K* different states the brain is in). For example, a simple model of brain connectivity would be that contralateral connections have some probability, and ipsilateral connections have a different one. One can further generalize this to a hierarchical block model, where each node is in a given set of nested groups [84–87]. For example, a node might be in a lobe within a hemisphere.

A further generalization asserts that each node is in its own group, and therefore has a “latent position” that characterizes its probability of connecting with other nodes (homologous to latent variable models in neural coding) [88]. A particularly popular version of these models assumes that the probability of connections between a pair of nodes is equal to the dot product between the nodes’ latent positions [89–91]. In these models, an extensive set of theoretical investigations have established the kinds of claims we desire when using a statistical model to make inferences about our data [92,93], as well as a number of extensions, including a generalized random dot product [94], a random dot product with node-wise covariates [95], and a latent structure model [96] (for review, see [97]). However, these models typically only operate on single, unweighted networks lacking attributes.

Some of these single-network models have been generalized to population models. One of the first such models was a non-parametric Bayesian model of populations of networks [98]. This model is essentially a network generalization of mixed effects models popular in biostatistics [99], where the mean network is a fixed effect, and each individual has a unique low-rank distortion relative to the mean. Two extensions of this approach are essentially non-Bayesian variants that enable faster computation [100,101], which was generalized to a mixture of random dot product models [102]. Estimation in each of these models can be thought of as specific tensor factorizations [103]. Although these models still lack much neuroscientific insight or attributes, they establish the statistical foundation for learning connectal codes.

## Statistical Model for Connectal Coding

To formalize connectal coding, we introduce the *connectal coding model*, which is designed to enable investigation of the links among connectypes, individual histories and cognitive phenotypes. Like all models, the connectal coding model makes simplifying assumptions for tractability. Although statisticians like to remind practitioners of George Box’s quip, “all models are wrong; some are useful”, we find this aphorism to be misleading. Models are maps, designed to get us from one place to another. The question is not “is a given map right or wrong?” but rather, “is the map useful for getting us to our destination?” The connectome coding model is designed to get us to a deeper understanding of the relationship among connectotypes, individual histories, and cognitive phenotypes. Insofar as it does that, it is useful.

In statistics, a code is a conditional distribution characterizing the probability of one random variable taking some value given another random variable taking some value. Let *X* and *Y* be random variables; their marginal distributions, *P[X]* and *P[Y]*, characterize the probability of any particular x or y, and their joint distribution *P[X,Y]* characterizes the probability of observing x and y. The conditional distribution *P[Y*|*X]* characterizes the probability that *Y* takes a particular value, given that *X* is some value. In connectal coding, we have the following four random variables:

- *B* = cognitive phenotypes of an individual, including and as measured by behaviors,
- *C* = connectome of an individual, spanning spatial and temporal scales,
- *D* = developmental history of an individual, including past experiences
- *E* = the current environment acting on individuals, and
- *G* = genome of an individual, including epigenetic factors.

Connectal coding concerns estimating the statistical relationships among connectomes and genomes, developmental histories, cognitive phenotypes, and the current environment. More formally, in connectal coding, we may seek to estimate the probability of a connectotype, given a genotype and environment, *P[C* | *D*, *E]*, and the probability of a cognitive phenotype, given a connectype and environment, *P[B* | *C, E]*. We are also interested in *P[C* | *D], P[B* | *C]*, and other conditional and joint distributions. In all cases, there exists a random variable that models the connectome, which therefore warrants further study. One random variable which may appear missing to many neuroscientists is brain activity. Much like one does not require modeling a connectome to characterize the relationship between stimuli/behaviors and brain activity, one does not require modeling activity to characterize the relationship between cognitive phenotypes/developmental history/genomes and brain connectivity. Although joint modeling of brain activity and connectivity would be more comprehensive, connectal coding is sufficiently challenging and interesting to warrant its own investigations.

This formalization of connectal coding, is, to our knowledge, novel. That said, this conceptual and formal model can be used to interpret previous connectomics studies, which often have similar conceptual frameworks [104]. More importantly, we hope it will help guide the development and ideation of new connectomic studies, by providing a paradigm within which to ask, formalize, and eventually answer questions about neural circuit mechanisms, how they work, how they fail, and how they can be improved.

### Connectal Coding Theories

Equipped with the statistical formalization of connectal coding, two questions we can ask are: (1) to what extent is a connectome statistically associated with “*X*” (e.g., genotype or cognitive phenotype), and (2) where in the connectome is that statistical association (that is, what is the connectotype associated with that genotype/phenotype)? An example of the first question is, “if the genomes of two individuals differ, to what extent are their connectomes different?” An example of the second question is: “where in the connectome are those differences?” Causal questions, such as which connectotype is part of the implementation-level causal mechanism of a given cognitive phenotype, are also possible, but beyond the scope of this article because they are much more difficult to answer convincingly (but see [104] for a wonderful article on this topic focused largely on fMRI based connectome estimates).

Formally asking these questions requires assuming a statistical model. The first kind of question is essentially a hypothesis-testing question. The data required to answer it are two collections of estimated connectomes: *C_1_,…C_n_* are estimated connectomes from individuals with one property (e.g., genotype or cognitive phenotype), and *C_n+1_,…, C_n+m_* are estimated connectomes from individuals with another property. Assume that *C_1_,…C_n_* are all sampled independently and identically from some distribution *P_0_ = *P[C* | *X* = 0]*, and *C_n+1_,…,C_n+m_* are sampled independently and identically from another distribution *P_1_* = *P[C* | *X* = *1]*. Then, the formal statement of the hypothesis is:

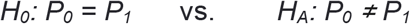

Applying such a test requires defining a test statistic, which quantifies the effect size, and computing the distribution of the test statistic under the null. Such two-sample tests have been developed for random graphs [105–108]. The theoretical claims associated with each of these two-sample testing results depends on an underlying statistical model of random graphs, such as those described in the previous section. These tests are holistic: they tell the researchers that there are differences between these populations and the magnitude of those differences (the test statistic), but they do not indicate where those differences are.

To answer the second question, global graph features (such as modularity) are inadequate because they cannot indicate *where* the differences are. Moreover, edge-wise statistics are typically underpowered when suitably adjusted for multiple comparisons, given our small sample sizes. Instead, we can search for a “signal subgraph”, that is, a small set of nodes and edges among them that confer the majority of the signal (the signal subgraph is an estimate of the connectotype). Signal subgraph searches are akin to feature screening, where one seeks to determine which features are most informative about a specific covariate [109]. The main difference is that signal subgraph methods take advantage of the graph structure to improve their sensitivity and specificity [110]. The first signal subgraph method used a variant of a sparse inhomogeneous Erdos-Renyi random graph model [111]. These methods have been extended to operate under latent variable model assumptions [112], and also to deal with continuous (rather than categorical) covariates [113]. Formally, signal subgraph detection is an estimation, rather than testing, problem: it seeks to estimate the smallest set of nodes such that the covariate is independent of the remaining nodes, and the signal subgraph is the set of edges among those nodes that carry information about the covariate.

## Applications

Consider the connectome of the larval *Drosophila* mushroom body [50] (Figure 1B). Priebe et al. [51] conducted an extensive empirical investigation of this connectome, leveraging spectral modeling, which resulted in the development of the latent structure random graph model [96]. Specifically, they discovered that kenyon cells of the mushroom body form a one-dimensional submanifold in a six-dimensional latent space. Figure 2 shows the quality of fit of several of these models to a binary simplification of the left larval mushroom body *Drosophila* data. These models provide a foundation from which to formulate statistical tests to answer the connecal coding questions described above.

**Figure 2:**
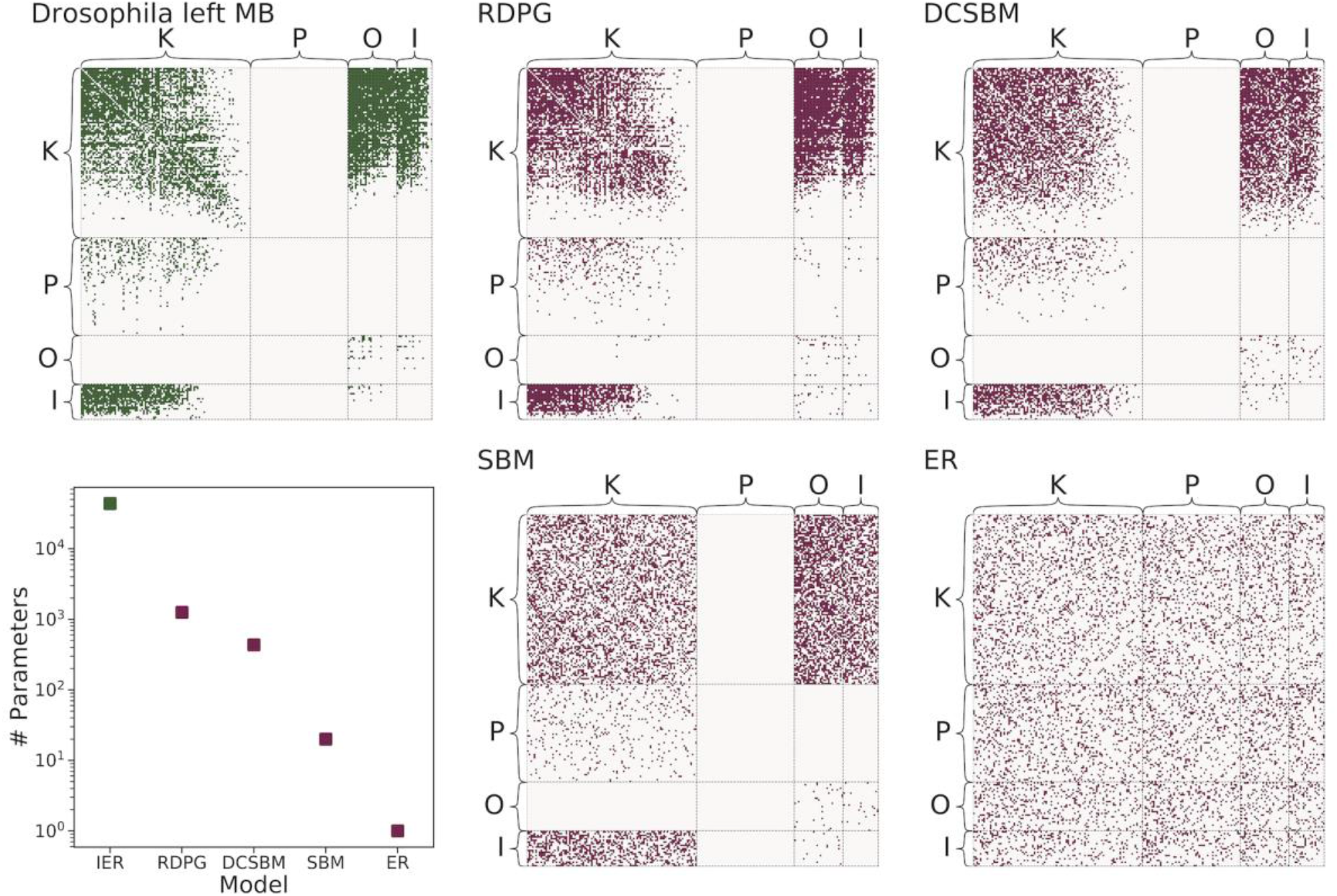
Connectome model fitting and complexity. Left larval *Drosophila* mushroom body adjacency matrix, followed by random samples from four different statistical models of connectomes with decreasing complexity: inhomogeneous Erdos-Renyi (IER), random dot product graph (RDPG), degree-corrected stochastic block model (DCSBM), stochastic block model (SBM), and Erdos-Renyi (ER). The bottom left shows the number of parameters for each. All graphs are sorted by node degree within each block.

Second, consider sex differences in the mouse brain estimated via diffusion magnetic resonance imaging (Fig. 1C). From scans of 55 mouse brains, 32 male and 23 female, connectomes were estimated on 332 nodes, 166 per hemisphere, using the Waxholm atlas [114]. The signal subgraph method of Wang et al. [112] reveals that 10 of the original 332 nodes contain more signal than noise about sex. The top-ranked nodes include a thalamic component and the periaqueductal gray, both of which are important in sexually dimorphic mouse brain development [115,116].

Third, we consider the COBRE data set [117], a collection of 123 functional MRI scans of schizophrenic and healthy human patients. Each scan yields an estimated connectome with 264 nodes, corresponding to 264 brain regions of the Power parcellation [118], with edge weights given by correlations between BOLD signals measured in those regions. The data set contains scans for 54 schizophrenic patients and 69 healthy controls (like the one shown in Figure 1D). Levin at al. [119] apply their omnibus embedding to jointly estimate a random dot product graph for each connectome, resulting in an estimated three-dimensional latent position for each node of each connectome. For each node, Hotelling’s T^2^ test yields a p-value assessing whether or not the latent positions from healthy connectomes are drawn from the same (normal) distribution as the latent positions from the schizophrenic connectomes. Because the Power parcellation is hierarchical, the nodes can be further organized into 14 parcels. Visualization of the distribution of these p-values within each parcel suggests that the connectomes of schizophrenics differ from those of healthy controls in certain subnetworks but not others (Fig. 3). Specifically, nodes in the default mode network tend to have significantly smaller p-values than nodes in, for example, the visual subnetwork. Independent investigations also implicate the default mode in schizophrenia [120,121], suggesting that our framework can provide statistical rigor to support previous scientific claims.

**Figure 3:**
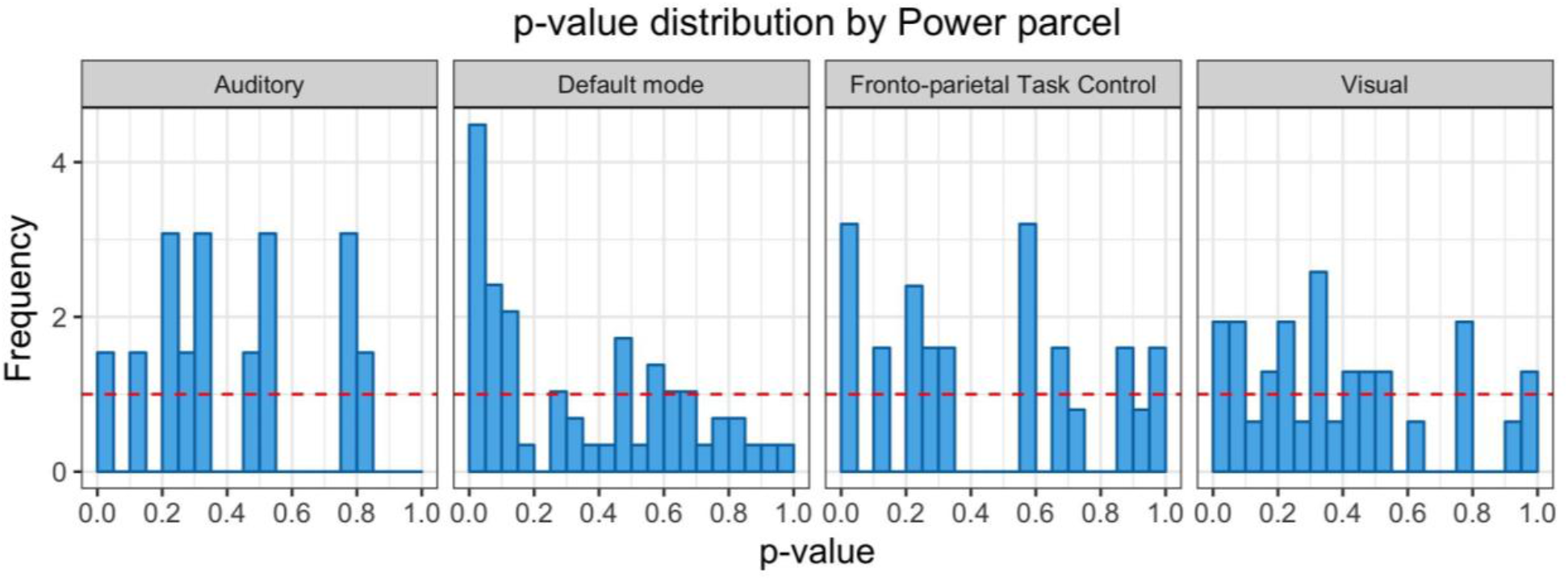
Normalized histograms of the distribution of p-values obtained from applying Hotelling’s T^2^ test to the omnibus embeddings of brain regions from a few selected Power parcels. The red dashed lines indicate the uniform density, which would be expected to hold if there were no difference between healthy and schizophrenic patients. The default mode network clearly displays a non-uniform p-value distribution, suggesting that this parcel differs in schizophrenic patients compared to their healthy counterparts. In contrast, p-values in the auditory, visual, and front-oparietal task control subnetworks appear approximately uniform, providing weak evidence that these systems are not implicated in schizophrenia.

## Discussion

Here, we discuss a few additional applications of the connectal coding framework. First, one can use connectal coding to study cognitive disorders, such as schizophrenia, as described above. The consensus report of the American Psychiatry Association’s working group on neuroimaging markers of psychiatric disorders concluded that “there are currently no brain imaging biomarkers that are currently clinically useful for any diagnostic category in psychiatry.” [122]. This is despite nearly 30 years (at the time of that report) of brain imaging. A widely held belief is that psychiatric illnesses are disorders of neural circuitry, or connectopathies [123–128]. If true, our ability to develop clinically useful prognostic, diagnostic, and treatment protocols will depend on connectal coding. The same strategy could be applied to study healthy brains as well. For example, memories are believed to be stored in “engrams”, which are defined as biophysical or biochemical changes in the brain underlying memories [129]. While engrams for specific memories largely remain elusive [130], connectal coding could accelerate this search, by formulating the specific memory as a cognitive phenotype. In these settings, heterogeneity of connectomes may be a significant impediment which would require further methodological developments. Further, the methodologies developed to study contrasts within and across individuals within a species can also be applied across species [131,132], although such studies should also take into account differences in body plan and life cycle.

Second, in other fields of inquiry such as particle physics and geology, simulations play a key role in understanding nonlinear dynamical systems. In brain science, however, simulations remain in their infancy. That said, recently, several high-profile efforts have emerged to simulate the brain of various species, including humans [133–136]. While the precise detailed requirements for biofidelic and useful simulations of a brain remain fiercely debated [137], it is unambiguous that *some* assumptions about nodes, their properties, and connections are required for any such simulation. Therefore, connectal coding could be exploited to learn which connectotypes are required for the simulation to exhibit which cognitive phenotypes.

Third, as alluded to above, the idea that understanding biological intelligence can inform machine intelligence dates back to the early days of computer science and the so-called first wave of artificial intelligence (AI). The first serious artificial neural networks were simple one-layer networks, called perceptrons [138]. The second wave of AI began when a few people realized that artificial neural networks, like biological neural networks, could have multiple layers to increase their expressive capacity. More formally, while perceptrons (one-layer networks) can only represent linear functions [139], multi-layer perceptrons (with only one hidden layer) can represent arbitrarily complex functions [140,141]. The third wave artificial intelligence, brought about by deep learning [142], appreciated that real connectomes do not have huge unstructured single hidden layers, but rather, many relatively small hidden layers [143,144]. Perhaps incorporating more biological constraints in these searches could further improve their efficiency [145]. Indeed, there are some signs that the third wave of artificial neural networks might be waning and/or asymptoting, and some posit that their main hope for continued rapid improvement is to incorporate more ideas from connectomics [146].

Connectal coding is an approach to model brain structures as networks and apply the wealth of statistical pattern recognition techniques to relate individuals’ networks to their phenotypes. To facilitate using these ideas, we have developed an open source python toolbox for statistical analysis of populations of networks, available at https://neurodata.io/graspy/ [149].

## Acknowledgements

The authors from JHU acknowledge support from NSF 16-569 NeuroNex contract 1707298, and KL’s support comes from NSF grant DMS-1646108. The authors are grateful for helpful feedback from Konrad Koerding. The authors declare no conflicts of interest.

## Footnotes

1 We acknowledge that it is common in the neuroscience community to ascribe other meanings to the word code, including representational and causal meanings [57]. We use “code” the way Shannon and information theory uses it, purely as a statistical relationship.

